# *Arabidopsis* CPK6 regulates drought tolerance under high nitrogen by the phosphorylation of NRT1.1

**DOI:** 10.1101/2022.10.06.511240

**Authors:** Qijun Ma, Chunyan Zhao, Shi Hu, Kaijing Zuo

## Abstract

Nitrogen is an essential macronutrient for plant growth and development, and its availability to some extent is regulated by drought stress. Calcium-dependent protein kinases (CPKs) are a unique family of Ca^2+^ sensors with diverse functions in nitrogen and drought signaling pathways. However, which and how CPKs involve in the crosstalk between drought stress and nitrogen transportation remains largely unknown. Here we identified the drought tolerant function of *Arabidopsis* CPK6 under high nitrogen condition. The *CPK6* expression is induced by the treatments of ABA and drought. The mutant *cpk6* is insensitive to the ABA treatment, but sensitive to drought only under high nitrogen condition. CPK6 interacts with and phosphorylates the Thr571 in NRT1.1 protein, and thus represses its NO_3_^−^ transporting activity under drought stress. Taken together, we showed the evidences that CPK6 regulates *Arabidopsis* drought tolerance through the phosphorylation of NRT1.1, and that enriches the knowledge of nitrogen uptake in plants during drought stress.

## Introduction

Nitrogen (N) is a macronutrient element sustaining plant growth and development (Krapp *et al.*, 2015; Vidal *et al.*, 2020). The absorption and utilization of nitrogen is a complicated process delicately regulated by various transcription factors and kinases under different environments such as drought stress (Gaudinier *et al.*, 2018; Brooks *et al.*, 2019; Vidal *et al.*, 2020). Nitrogen metabolism (absorption and utilization of nitrogen) and drought stress (water deficiency) are interconnected physiological processes in plants, and there are molecular links between two signaling pathways (Plett *et al.*, 2020; Shangguan *et al.*, 2000).

Drought stress reduces N absorption or/and assimilation of plants through down-regulating the enzyme activities of nitrate reductase (NR), glutamine synthase (GS) and glutamate synthase (GOGAT) in roots and leaves (James *et al.*, 2018; Iqbal *et al.*, 2020; Han *et al.*, 2022). Drought stress also has great impacts on the activities of nitrogen transporters. In the *NRT1.1* knockout *Arabidopsis* mutant, the decreased root hydraulic conductivity was correlated with shoot NO_3_^−^ concentration (Guo *et al.*, 2003). Polyethylene glycol (PEG) treatment upregulated the expression of the *NAR2* genes (*NRT3* genes) and decreased the expression levels of several *NRT2* genes (Cao *et al.*, 2018). Overexpression of *OsNAR2.1* raised the vegetative growth of rice under the PEG treatment with higher grain yield (Chen *et al.*, 2019). Besides above N transporters, the stability of NRT1.2 and its ABA transporting activity are precisely regulated by CEPR2 in response to environmental stress such as drought (Li *et al.*, 2016; Kamiya *et al.*, 2012; Zhang *et al.*, 2021). Recently, two wheat varieties overexpressed a drought-responsive transcription factor *GmTDN1* have enhanced N absorption by upregulating expressions of *NRT2.5* (Zhou *et al.*, 2022), strongly suggesting that drought stress influences different components of N transportation and assimilation systems.

From another aspect, the levels of N availability reversely regulate drought tolerance through mediating root water uptake and photosynthesis (Hepworth *et al.*, 2015; Song *et al.*, 2019a, b). Low soil nitrogen (low-nitrogen, LN) generally enhances plant photosynthesis for adaptation to water-deficit stress (Hepworth *et al.*, 2015). The pretreatment LN on plants has the potential to mitigate the adverse effects of abiotic stresses (Gao *et al.*, 2018). Under high nitrogen (HN) condition, maize plants are more sensitive to drought stress with reduced stomatal closure and leaf elongation rates (Xing *et al.*, 2021). These results further indicate that there are tight crosstalks between drought stress and N metabolism signalling, and the signalling integrators remain to be explored (Han *et al.*, 2022).

NRT2.1 and NRT1.1 are considered as two crucial transporters of plants in most of growth environments (Wang *et al.*, 2012; Vidal *et al.*, 2020). AtNRT2.1 could be rapidly dephosphorylated with the addition of HN but remained phosphorylated state under LN condition (Engelsberger *et al.*, 2012; Ohkubo *et al.*, 2021). The dephosphorylation of AtNRT2.1 is important for limiting root N uptake in response to HN supply (Filleur *et al.*, 2001; Engelsberger *et al.*, 2012; Ohkubo *et al.*, 2021). Under HN condition, drought stress had 30% reduction in root hydraulic conductivity of *NRT2.1* mutant plants (Li *et al.*, 2016). In response to environmental nitrate change, NRT1.1 in *Arabidopsis* functions as a dual affinity transceptor involving both high- and low-affinity uptake through the phosphorylation/dephosphorylation of Thr^101^ (Ho *et al.*, 2009). Under LN condition, CIPK23 phosphorylates the Thr^101^ in AtNRT1.1 serving as a high-affinity nitrate transporter (Ho *et al.*, 2009; Maghiaoui *et al.*, 2020). Upon HN supply, NRT1.1 in *Arabidopsis* is dephosphorylated by the phosphatase ABI2 and transits into a low-affinity nitrate transporter (Ho *et al.*, 2009; Sun *et al.*, 2014; Léran *et al.*, 2015; Maghiaoui *et al.*, 2020; Li *et al.*, 2020). Due to the importance of biological functions in drought tolerance and N transporter, NRT1.1 probably acts as a bridge connecting the signaling pathways of drought stress and N transportation (Guo *et al.*, 2003).

CPKs is a kind of calcium sensors composed of a variable N-terminal domain, a Ser/Thr protein kinase domain and an auto-inhibitory junction domain, which mediate various physiological processes and biotic/abiotic stress responses. CPKs function in multiple signal transduction pathways by phosphorylating various ion channels, transcription factors and metabolic enzymes during plant growth, development and the tolerance to biotic/abiotic stresses (Yip and Boudsocq, 2019). CPK10, 30, 32, the master regulators of nitrate signalling, phosphorylate the NIN-LIKE PROTEIN (NLP) transcription factors to coordinate plant growth and development (Liu *et al.*, 2017). CPK32 interacts with AMMONIUM TRANSPORTER 1;1 (AMT1;1) to positively regulate ammonium uptake in roots (Qin *et al.*, 2020). Besides them, CPK6 has important functions in biotic and abiotic stress through phosphorylating different substrates of SLAC1 and SLAH3. For example, CPK6 participates in ABA regulation of guard cell through phosphorylating SLAC1 for stomatal closure (Mori *et al.*, 2006; Brandt *et al.*, 2012). CPK6 mediates plant drought tolerance by interacting with and phosphorylating transcription factors ABF3 and ABI5 in the ABA signaling pathway (Zhang *et al.*, 2020). These results together reveal that CPKs including CPK6 have emerged as the target in regulating nitrate transport and drought tolerance, but how do CPKs work in their crosstalk remains elusive. In this study, we uncovered the role of *Arabidopsis* CPK6 in drought stress by modulating the phosphorylation level of NRT1.1. We also provided evidences supporting CPK6 as a negative regulator of the nitrate transporter NRT1.1 under HN treatment. Hence, this study provides the clues for deeper understanding how plants uptake N from soil facing drought challenges.

## Materials and methods

### Plant materials and growth conditions

The seeds of T-DNA insertion mutants (*cpk6*, SALK_025460C; *chl1-5*, CS6384) were ordered from the AraShare center (Beijing, China). *Arabidopsis thaliana* ecotype (Columbia, Col-0) and the mutants were grown under the 16 h/8 h (light/dark) photoperiod at approximately 600 mmol m^−2^.s^−1^ at 25℃. To analysis ABA sensitivity, the seeds of Col-0, *cpk6* were treated with 2% NaClO_3_ for three times (5 min for each treatment), and washed with sterile water for five times (each wash lasted 5 min). The sterilized seeds were sowed on the medium with 0 μM and 0.1 μM ABA combined with 1 mM or 10 mM KNO_3_ respectively, and then incubated at 4 ℃ for 2 d. After that, all the seedlings were grown under the 16 h/8 h (light/dark) photoperiod at approximately 600 mmol. m^−2^.s^−1^ at 25℃.

For the drought tolerance assays, the sterilized seeds (Col-0, *cpk6*, *chl1-5* and *cpk6/chl1-5*) were firstly germinated on 1/2 MS medium for 7 days. The seedlings were then transferred into soil without watering for 15 d creating drought stress. Each treatment contained at least 10 pots with at least 100 seedlings. The experiments were repeated at least three times. Statistical analyses of the numerical data were conducted by Microsoft Excel (2003) and SPSS statistical software 13.0 (SPSS Inc.).

### Cloning full-length cDNA of CPK6 and NRT1.1 genes

Total RNAs were extracted from the leaf tissues of Col-0 plants using Trizol Reagent (Invitrogen Life Technologies, Carlsbad, CA, USA). DNA contamination was removed with an on-column DNase treatment followed the kit menu (Thermo-Fisher, USA). Total RNA (1mg) was reverse-transcribed into the first-strand cDNA in a 20 μl reaction volume using the first strand cDNA synthesis Kit (TaKaRa, Dalian, China). PCR amplification were 94℃ for 3 min; followed by 30 cycles of 94℃ for 30 s, 56℃ for 40 s and 72℃ for 2 min; with a final extension at 72℃ for 10 min. The amplified fragments were ligated into vector *pDONR* and sequenced. Primers were presented in the **Table S1**.

### Quantitative real-time RT-PCR (qRT-PCR) analysis

The leaves of Col-0, mutant plants under normal condition and drought stress were harvested respectively. All the leaves were used to extract the total RNAs immediately; To analysis the expression profiles of *CPK6* under the ABA treatment, 12-days *Arabidopsis* seedlings grown in the medium with 1mM or 10mM nitrate respectively were treated with 0, 0.1 μM ABA. Whole seedlings were collected to extract the total RNAs.

Total RNAs (1mg) were reversely transcribed into the first-strand cDNA according to the method above described. qRT-PCR analysis were performed with iQ-SYBR Green Supermix kit in an iCycler iQ5 system following the instructions of the manufacturer (Bio-Rad, USA). The relative quantifications of mRNA levels were performed using the cycle threshold (Ct) 2^−∆∆Ct^ method.

The expression levels of *CPK6*, *LBD39, LBD38, MYC1, NRT1.1, NRT2.1, NRT2.5, HAB1, RD29A, COR15, DREB2A, RD22, COR47* and *EXLB1* in *Arabidopsis* were analyzed with the primers listed in Table S1. The gene expression levels were normalized against the signals of *Actin 7* gene. The gene *Actin* 7 was amplified with primers ACTIN 7-1 and ACTIN 7-2, and used as the internal reference. All of the samples were tested at least three biological replicates.

### BiFC analysis

The full-length cDNA of *CPK6* and *NRT1.1* gene without the termination codon were cloned into *pEarleyGate202* and *pEarleyGate20* vector respectively. The positive plasmids were co-transformed into the *Agrobacterium* strains *EHA105* using chemical transformation. The p19 protein of tomato bushy stunt virus was used to suppress gene silencing. For co-infiltration, equal volume suspensions of different *Agrobacterium* strains carrying different constructs *pEarleyGate201* and *pEarleyGate202* were mixed with the ratio 1:1 prior to infiltration. The resuspended agrobacterium cells were infiltrated into leaves of tobacco plants as described previously (Ma *et al.*, 2019). Three days later, the YFP fluorescence signals showing the protein interaction between NRT1.1 and CPK6 were visualized under the confocal laser scanning microscopy and digitally recorded (TCS SP5, Leica, Germany).

### Protein extraction and western blotting

A quantity of 2 g of *Arabidopsis* leaves for each sample was ground into the fine powder and dissolved into the buffer containing: 20 mM Tris-HCl (pH 7.4), 100 mM NaCl, 0.5% Nonidet P-40 (v/v), 0.5 mM EDTA, 0.5 mM PMSF, and 0.5% (v/v) Protease Inhibitor Cocktail (Sigma-Aldrich). The total proteins were extracted according to the reported method (Ma et al., 2019). After demining the concentration of total protein, 10 μg total proteins were separated on a 12% SDS-PAGE gel and transferred to the PVDF membranes (Roche, CA, USA) using an electro-transfer apparatus (Bio-Rad). The membranes were incubated with the GFP or NRT1.1 (Sigma-Aldrich, USA) primary mono-antibodies, and then hybridized with the peroxidase-conjugated secondary antibodies (Abcam, Shanghai, China). The western blotting signals were visualized using an ECL Detection Kit (Millipore). The actin protein served as the protein loading control.

### The split-ubiquitin membrane-based yeast two-hybridization

The DUAL membrane system based on the split-ubiquitin mechanism was used to detect the interaction between CPK6 and NRT1.1 (Ma *et al.*, 2019). CPK6 were respectively fused to the C-terminus of ubiquitin with an artificial transcription factor LexA-VP16 as the baits, meanwhile NRT1.1 was fused with the N-terminal part of ubiquitin as the prey. The interaction was determined by the growth of yeast colonies on the SD medium lacking -T/-L/-H/-A.

### In vitro pull-down assay

The coding sequence of *NRT1.1* gene with *BamHI* and *SalI* adaptors was digested with *BamHI* and *SalI* endonucleases, and cloned into p*YES2* vector and fused with the His-tag. The full-length cDNA of *CPK6* gene with adaptors was digested by *EcoRI* and *SalI* endonucleases and cloned into pGEX4T-3 vector fusion with the GST tag. The plasmids of pGEX-CPK6-His and p*YES2*-CPK6-GST were transformed into *Escherichia coli* BL21 (DE3; Transgene) and BY4741, respectively. For pull-down analysis, CPK6-GST proteins were firstly eluted from the glutathione-agarose beads, and then incubated with the NRT1.1-His fusion protein which remained attaching to the tetradentated-chelated nickel resin. In general, proteins were incubated at 4°C for at least 4 h under the gently shaking condition. After incubation, the precipitants were washed at least three times to remove unspecific bound proteins. The pull-down proteins were finally boiled at 100°C for 10 min. The proteins were run on the 12% the SDS-PAGE gel, and detected with His and GST antibodies (Sigma-Aldrich, USA), respectively.

### In vitro phosphorylation assay

To analysis the phosphorylation activities of CPK6 and NRT1.1, the fusion proteins NRT1.1-His and CPK6-GST were expressed in the tobacco epidermal cells according to the reported method (Ma et al., 2019). A total of 0.2 μg of recombinant protein NRT1.1-His and 1 μg of CPK6-GST fusion proteins was incubated in 25 μL of reaction buffer [20 mM Tris-HCl (pH 7.5), 5 mM MgCl_2_, 10 mM NaCl and 2 mM DTT] with y-s-ATP at room temperature for 30 min. The phosphorylated proteins were visualized using autoradiography after separation on a 12% SDS-PAGE gel.

### Protein phosphorylation analysis by mass spectrometry

Protein phosphorylation analysis by mass spectrometry was performed according to the reported method (Garcia et al., 2005). To identify the possible phosphorylation sites in NRT1.1 protein, the *Arabidopsis* protoplasts expressing the fusion protein NRT1.1-GFP and CPK6-Flag were generated. Western blot indicated that the slow-moving proteins were collected with anti-GFP antibody-conjugated agarose beads and separated in SDS-PAGE gel. After proteolytic digestion and purification, the protein samples were analyzed by the liquid chromatography-tandem mass spectrometry (LC-MS/MS).

### Measurement of the net NO^3−^ flux using the scanning ion-selective electrode technique

The seeds of Col-0, *cpk6* were sterilized according to the method describe above. The sterilized seeds were sowed on the medium without nitrogen and grew for 7 days under the 16 h/8 h (light/dark) photoperiod at approximately 600 mmol.m^−2^.s^−1^ at 25℃. After that, the seedlings were treated with different concentrations of nitrogen (^15^N) and 300 mM mannitol. After 5 min, the seedlings were collected and determined by LC-MS. The net fluxes of NO_3_^−^ were measured non-invasively using SIET (BIO-003A system; Younger USA Science and Technology Corp., Amherst, MA, USA; Applicable Electronics Inc., Forestdale, MA, USA; Science Wares Inc., Falmouth, MA, USA). The analytical method followed the previously reported and the instrument protocol (Sun *et al.*, 2014).

## Results

### The mutation of *CPK6* in *Arabidopsis* decreases drought tolerance under high nitrate condition

To analysis whether the drought tolerant function of *CPK6* is related to nitrogen concentration, we investigated the phenotypes of *cpk6* mutant and Col-0 plants grown in the soils with LN (1 mM) and HN (10 mM) respectively (Figure 1A). No significant difference was observed in the growth between Col-0 and *cpk6* mutant plants under LN condition after drought treatments (Figure 1A). Col-0 and *cpk6* mutant plants showed the identical phenotypes including fresh weight and the contents of malondialdehyde (MDA), and the growths (Fig. 1A-C). In contrast, the *cpk6* mutant plants were sensitive to the drought treatment under HN condition (Fig. 1a). The fresh weight was significantly reduced; and the MDA content was obviously increased in the *cpk6* mutant plants after drought treatment (Fig. 1B-C). The *cpk6* plants became more wilted and yellower than Col-0 plants after the drought treatment for 1 days (Figure 1A). The expressions of drought responsive genes (*RD29A, COR15, DREB2A, RD22, COR47, EXBL1*) were significantly decreased in *cpk6* mutant than Col-0 plants in the HN soil under drought stress (Figure 1D). These results indicated that the *cpk6* mutant is sensitive to drought stress under HN condition, and the drought tolerant function of *CPK6* is related to environmental nitrogen concentration.

**Figure 1.**
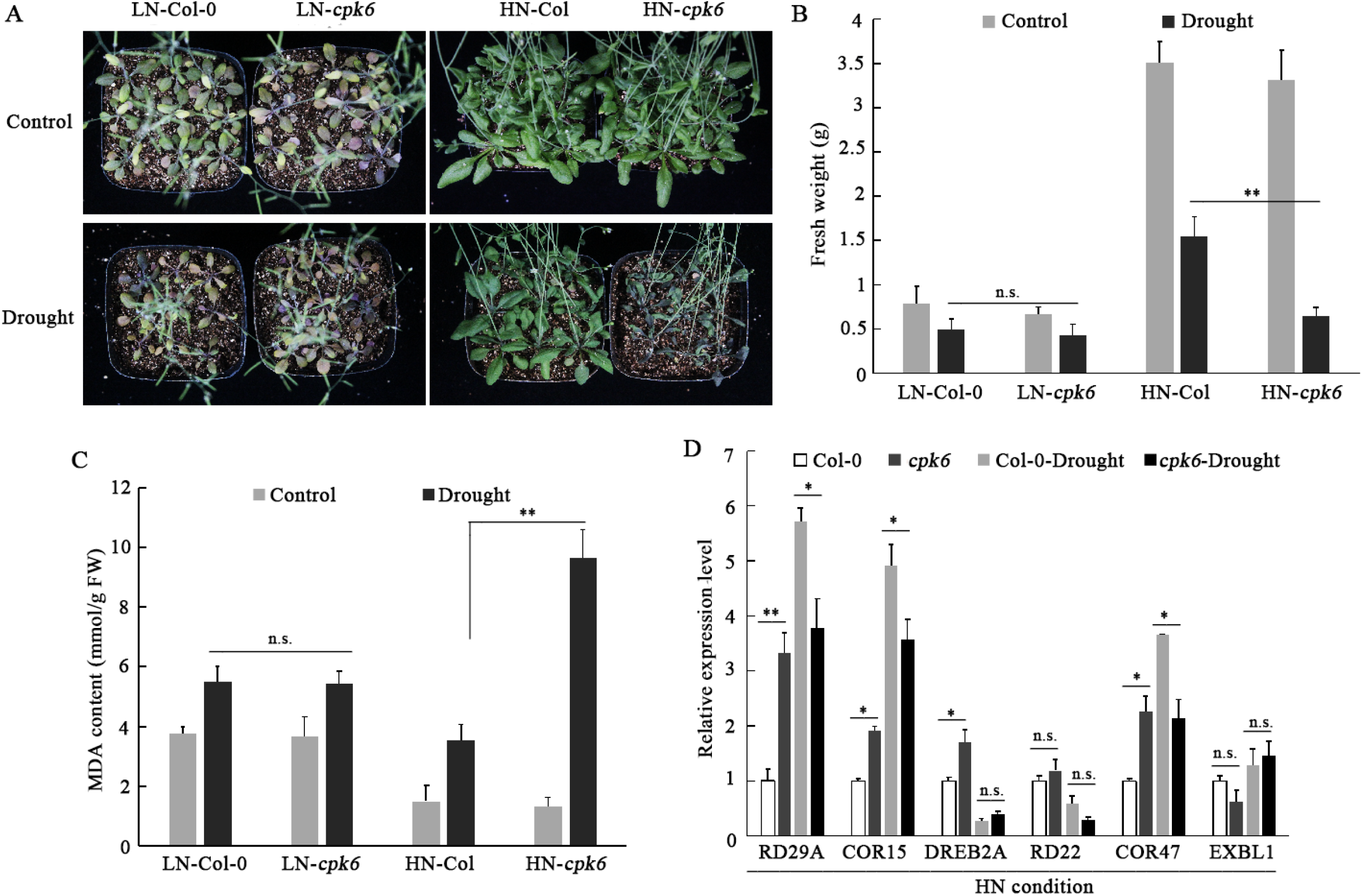
The *Arabidopsis* mutant *cpk6* confers sensitive to drought under high nitrogen condition. **(A)** The drought tolerance phenotypes of the *cpk6* mutant and Col-0. The *cpk6* mutant and Col-0 plants grown in the soil of LN (1mM KNO_3_) and HN (10mM KNO_3_) conditions were treated with drought for 15 days and then photographs were taken. **(B)** and **(C)** The fresh weight and MDA content in the mutant *cpk6* and Col-0 plants grown in LN and HN conditions treated with drought stress. Each column represents the mean±SD. of three independent replicates. **(D)** The relative expression levels of 6 drought stress related genes (*RD29A, COR15, DREB2A, RD22, COR47, EXBL1*) in the mutant *cpk6* and Col-0 plants under LN and HN conditions after the drought treatment. All experiments were performed in triplicate.

*CPK6* expression was repressed by the treatment of HN but increased by the treatment of both ABA and HN (Fig. 2A). We further investigated the growth difference between Col-0 and *cpk6* mutant seedlings grown in the 1/2MS medium with 2 μM ABA combined with different concentration of nitrogen. No significant difference in terms of root elongation was observed between the mutant *cpk6* and Col-0 seedlings grown in the medium with 1 mM NO_3_^−^ and 2 μM ABA medium. With the increased NO_3_^−^ in the medium, the *cpk6* seedlings became more insensitive to ABA than Col-0; and the *cpk6* seedlings had longer roots than *Col-0* in the medium of 3 mM and 10 mM NO_3_^−^ and 0.1 μM ABA (Fig. 2B-C), indicating that the ABA-insensitivity of *cpk6* is affected by nitrogen concentration.

**Figure 2.**
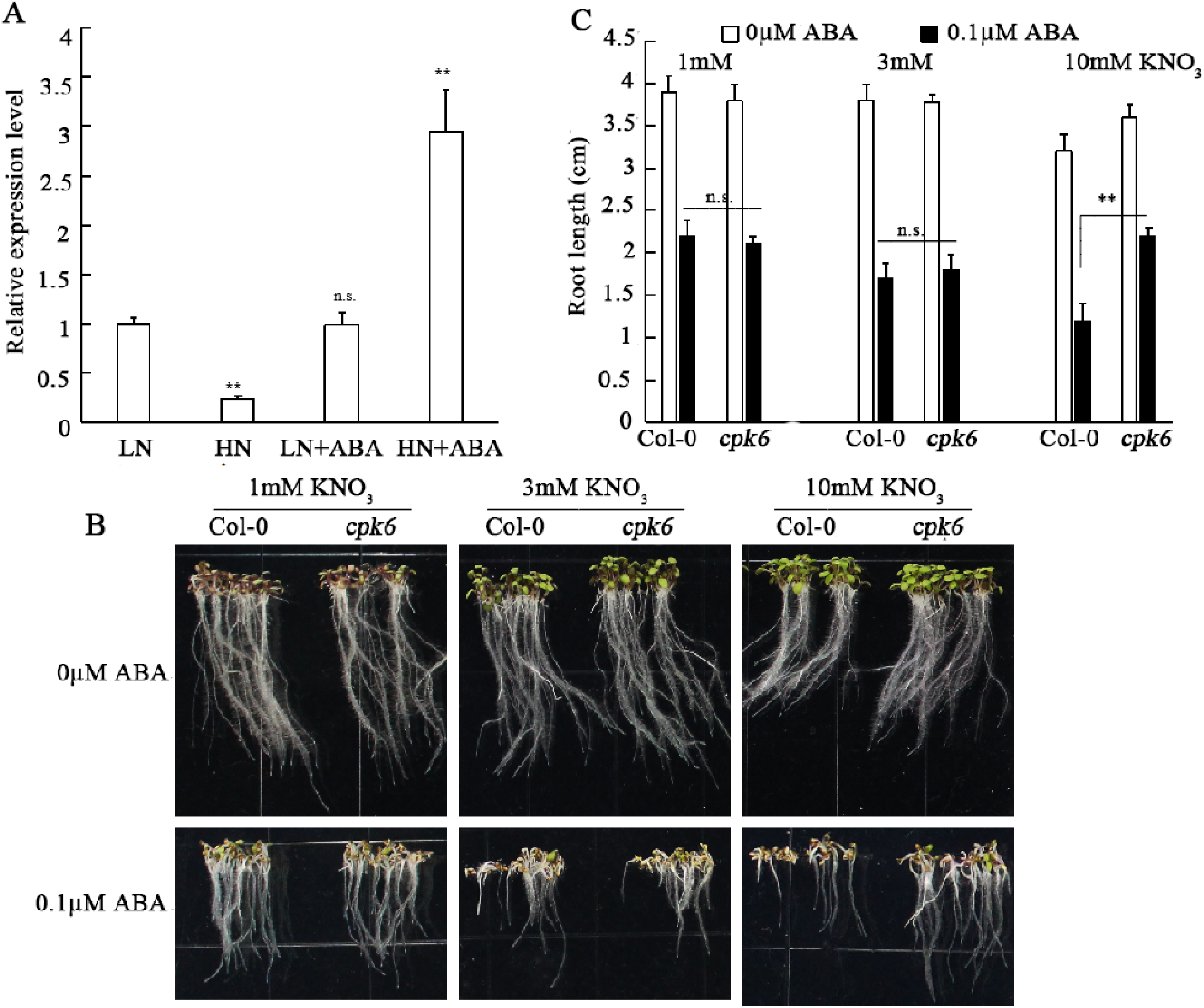
The *Arabidopsis cpk6* mutant loses ABA sensitively under HN condition. **(A)** Expression levels of *CPK6* gene in the *Arabidopsis* treated with LN (1mM KNO_3_), HN (10mM KNO_3_), LN (1mM KNO_3_)+0.1μM ABA and HN (10mM KNO_3_)+0.1μM ABA. **(B)** The seed of the mutant *cpk6* and Col-0 plants were grown in the medium containing 1, 3 and 10mM KNO_3_ or 1, 3 and 10mM KNO_3_+0.1μM ABA for 10 days and then photographs were taken. **(C)** The root length in the mutant *cpk6* and Col-0 plants grown in the medium containing 1, 3 and 10mM KNO_3_ or 1, 3 and 10mM KNO_3_+0.1μM ABA. Each column represents the mean±sd. of three independent replicates.

### The *cpk6* mutant reduces N absorption ability under drought treatment

Given that the drought tolerant function of *CPK6* is affected by HN, we further analyzed N absorption ability of *cpk6* mutant and Col-0 plants after drought treatment. The nitrate content of Col-0 and *cpk6* mutant plants had no significant difference when they grew under LN condition after drought treatments (Fig. 3A). However, the *cpk6* mutant plants had more nitrate content than Col-0 under HN condition after drought treatment (Fig. 3A). The mutant lines *cpk6* had significantly higher nitrate influx rates than those of Col plants in a solution containing 10 mM nitrate for 10 min (Fig. 3B). ^15^N absorption experiments results indicated that the drought treatment decreased N absorption abilities of both Col-0 and the *cpk6* mutant plants under HN. The decreased values of ^15^N absorption and the total content of NO_3_^−^ were less in the *cpk6* mutant than those of Col-0 (Fig. 3C). These results indicated that *cpk6* mutation increases the N absorption rate in *Arabidopsis* under drought condition and HN.

**Figure 3.**
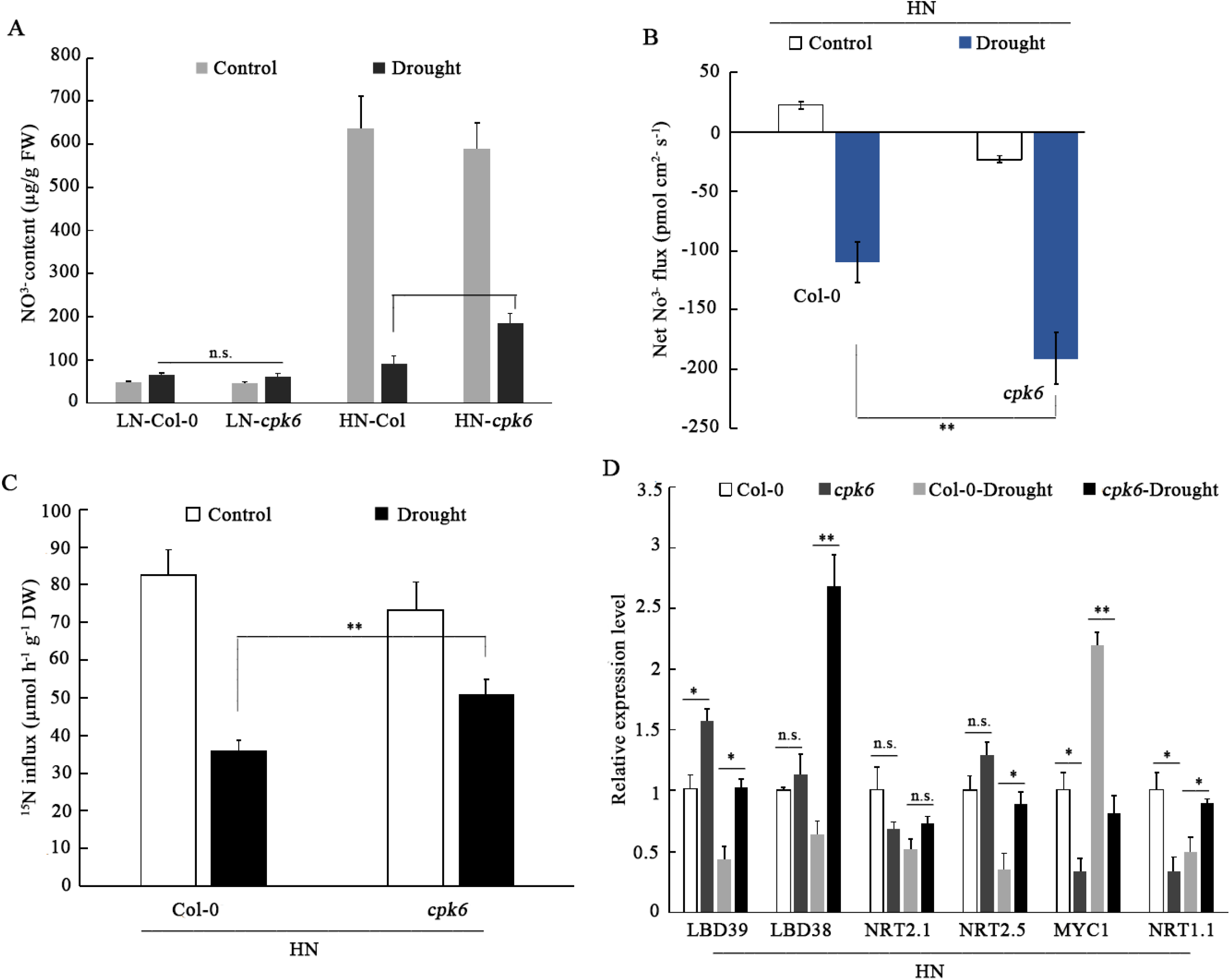
Knockout *CPK6* gene expression in Arabidopsis enhances N absorption abilityunder drought treatment. **(A)** The NO_3_^−^ content of *cpk6* mutant and Col-0 grown in the soil of LN (1mM KNO_3_) and HN (10mM KNO_3_) conditions treated with drought stress. **(B)** Net NO_3_^−^ flux of Col-0 and *cpk6* mutant with 10mM KNO_3_+drought conditions. **(C)** ^15^N influx of Col-0 and *cpk6* mutant grown in the soil of LN (1mM KNO_3_) and HN (10mM KNO_3_) conditions treated with drought stress. **(D)** The relative expressions of 6 nitrate related genes (*LBD39, LBD38, NRT2.1, NRT2.5, MYC1, NRT1.1*) in *cpk6* mutant and Col-0 grown in the soil of LN (1mM KNO_3_) and HN (10mM KNO_3_) conditions treated with drought stress. All experiments were performed in triplicate.

RT-qPCR showed that the expressions of nitrate signaling gene *LBD39* were significantly increased and MYC1 was repressed in both Col-0 and cpk6 plants under HN after drought treatment. The expressions of *NRT1.1* were changed conversely between *cpk6* mutant and Col-0 plants (Fig. 3D), indicating that knocked out *CPK6* expression influences N absorption ability and the expressions of nitrogen transporter genes under drought stress.

### CPK6 interacts with NRT1.1

Given that CPK6 is a kinase with phosphorylation activity, we used DUAL membrane yeast hybridization system to identify the interactor with CPK6 protein. After 2-round screening, a positive colony with the insert of *NRT1.1* gene was detected to be the CPK6 interaction candidate. Transient expression of mCherry-NRT1.1 and GFP-CPK6 fusion proteins in tobacco leaf epidermal cells confirmed that NRT1.1 co-localized with CPK6 in plasma membrane (Fig. S1). BiFC showed that the strong fluorescence signals were detected on the plasma membrane of tobacco leaf epidermal cells co-expressing cYFP-CPK6 and nYFP-NRT1.1, confirming the NRT1.1-CPK6 interaction in *vivo* (Fig. 4A). Yeast two hybridization assay was further conducted to validate the interaction between CPK6 and NRT1.1. Yeast cells containing the plasmids of pGAD-NRT1.1 plus pGBD-CPK6 grew well on the SD-Trp/-Leu/-His/-Ade medium (Figure S2). Meanwhile, the other CPK proteins including CPK5, CPK25 and CPK26 showed no interaction with NRT1.1 (Figure S2). Pull-down and co-immunoprecipitation (Co-IP) assays in tobacco leaf epidermal leaves also confirmed that NRT1.1 interacts with CPK6 in *vitro* and *vivo* (Fig. 4B and 4C).

**Figure 4.**
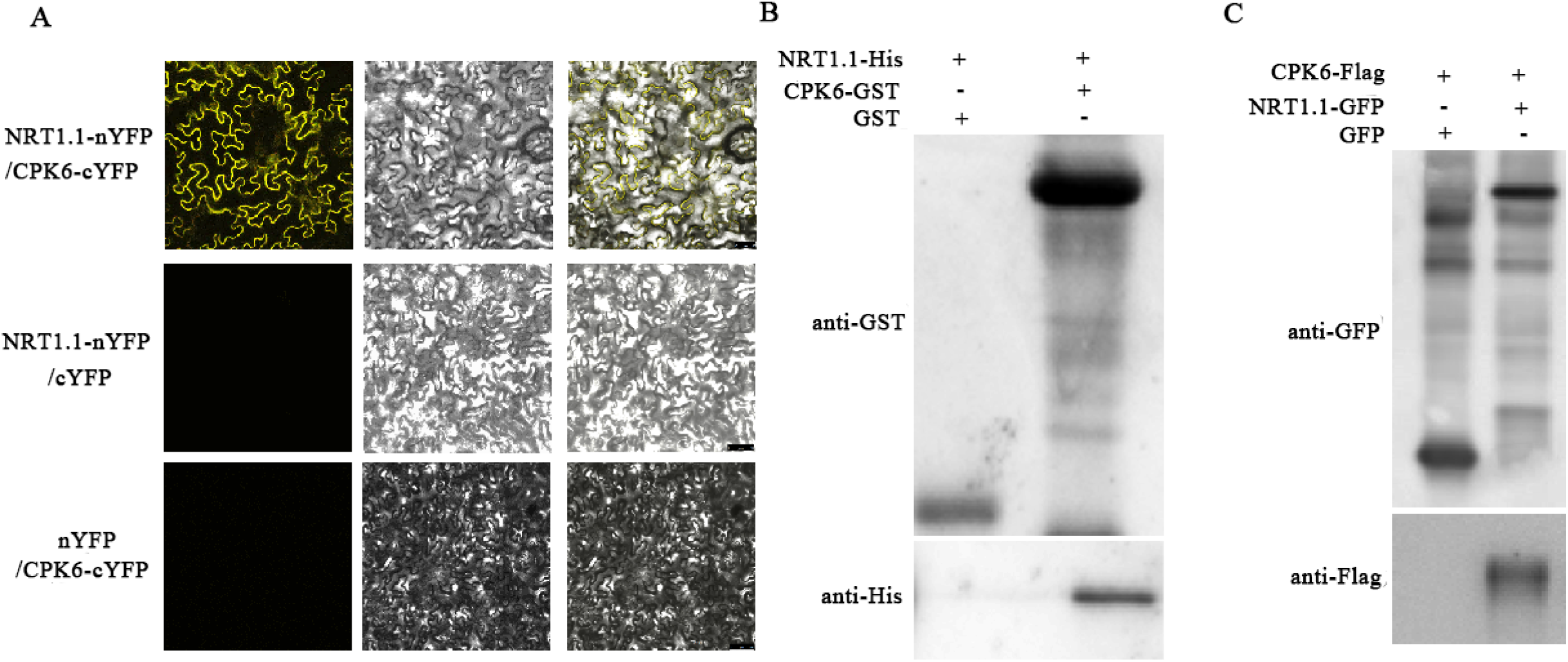
CPK6 interacts with NRT1.1 in *vivo* and in *vitro*. **(A)** Bimolecular fluorescence complementation *BiFC* analysis of the interaction between CPK6 and NRT1.1 in *N. benthamiana*. Scale bars represent 50μm. **(B)** Interaction between CPK6 and NRT1.1 proteins in *vitro*. NRT1.1-His proteins were incubated with immobilized CPK6-GST or GST, and the proteins immunoprecipitated with GST-beads were detected using anti-His antibody. **(c)** Interaction between CPK6 and NRT1.1 proteins in *vivo*. CPK6-Flag proteins were incubated with immobilized NRT1.1-GFP or GFP, and the proteins immunoprecipitated with GFP-beads were detected using anti-Flag antibody.

### Knocked out *CPK6* expression decreases the phosphorylation level of NRT1.1 in *Arabidopsis* under drought condition

CPK6 functions in the regulation of stomatal opening and drought tolerance under high nitrate condition (Guo *et al., 2003). Regarding that the interaction between CPK6 and NRT1.1, we further analyzed the p*hosphorylation level of NRT1.1 protein in *Arabidopsis* under LN and HN respectively using specific anti-NRT1.1 antibody. The phosphorylation levels of NRT1.1 protein was increased in both Col-0 and *cpk6* mutant under HN. However, the phosphorylation level of NRT1.1 in *cpk6* mutant under HN was significantly reduced after drought treatment compared with Col-0 (Fig. 5A-B). These results showed that *CPK6* expression change can influence the phosphorylation level of NRT1.1 in *Arabidopsis* especially under the treatment of HN and drought.

**Figure 5.**
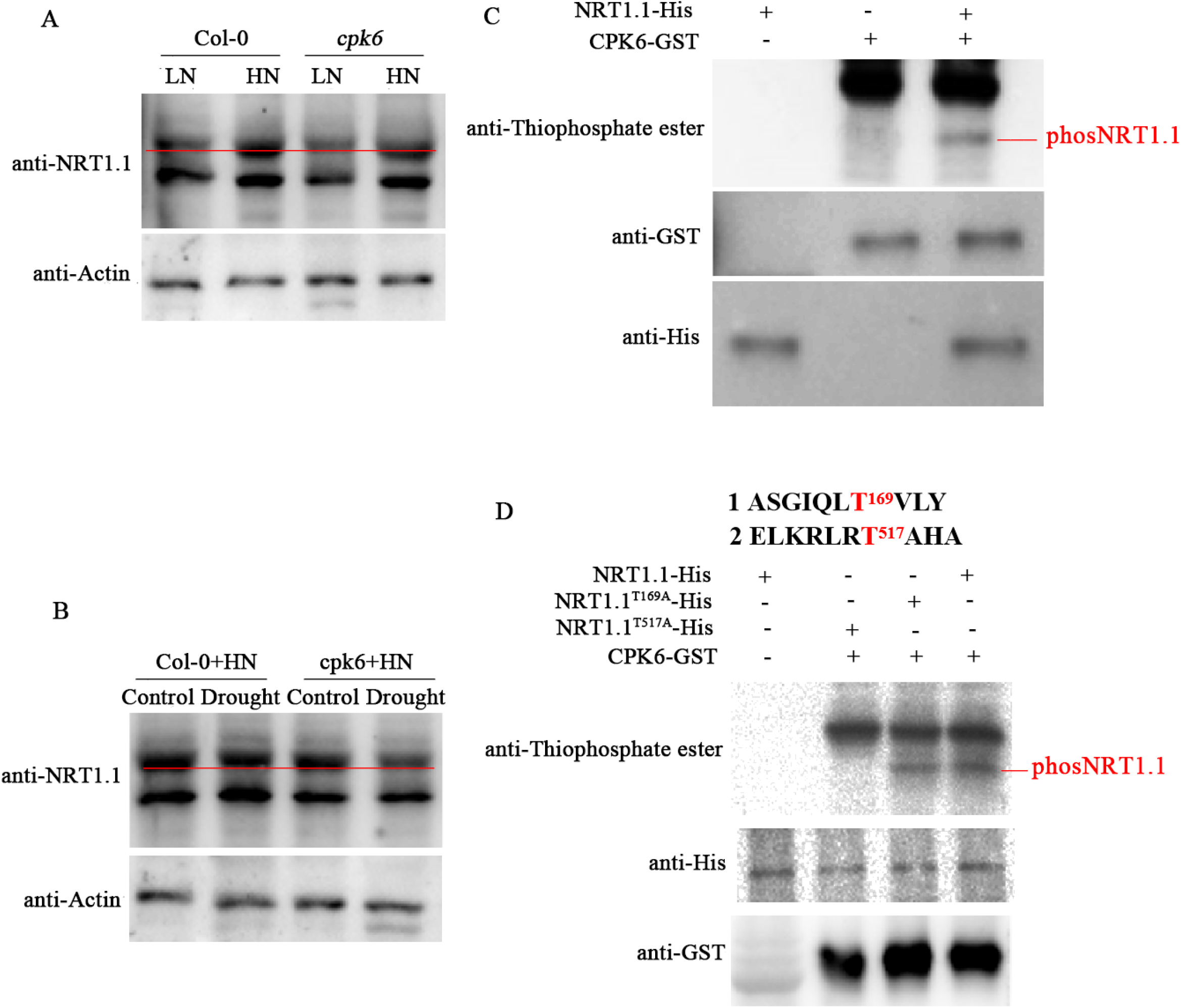
CPK6 mediates the phosphorylation of NRT1.1 in response to drought under HN conditions. **(A)** Western blot detected phosphorylation of NRT1.1 protein in the Col-0 and *cpk6* mutant plants using an anti-NRT1.1 antibody under LN and HN. **(B)** In *vitro* phosphorylation assay of NRT1.1 by protein kinase CPK6. SDS-PAGE gel with NRT1.1-His, CPK6-GST proteins (up panel); Thiophosphate ester showing NRT1.1 phosphorylation by CPK6. **(C)** Western blot detected phosphorylation level of NRT1.1 protein in the Col-0 and *cpk6* mutant plant using an anti-NRT1.1 antibody in response to drought stress under HN. **(D)** In vitro phosphorylation assay of NRT1.1 at Thr169 and Thr517 by protein kinase CPK6. SDS-PAGE gel blotted with anti-His and anti-GST antibodies detecting NRT1.1-His, NRT1.1^T169A^-His, NRT1.1^T517A^-His and CPK6-GST proteins (up panel). For western-blot assays, ACTIN protein was used as the loading control to ensure identity in the amounts of tested proteins.

To analysis whether CPK6 phosphorylates NRT1.1, we expressed NRT1.1-His and CPK6-GST protein and performed the phosphorylation assays in *vitro*. Western blotting showed that NRT1.1-His fusion protein was phosphorylated by CPK6-GST (Fig. 5C). To identify the phosphorylation sites of NRT1.1 protein, we digested and purified NRT1.1-His fusion protein and analyzed it by liquid chromatography-tandem mass spectrometry (LC-MS/MS). The result showed that the Thr^169^ and Thr^517^ in NRT1.1 protein were candidate sites phosphorylated by CPK6 under drought stress (Fig. S4; Fig. 5D). We further mutated these sites (NRT1.1^T169A^ and NRT1.1^T517A^) to mimic non-phosphorylated states and performed the phosphorylation assay. In *vitro* phosphorylation assays revealed that NRT1.1^T169A^ protein could still be phosphorylated by CPK6, whereas the NRT1.1^T517A^ lost phosphorylation activity (Fig. 5D). These results indicated that the Thr^517^ residue of NRT1.1 is phosphorylated by CPK6 protein.

### *CPK6* mutation decreases drought tolerance in *Arabidopsis* requiring NRT1.1 protein

To analysis whether the transportation activity of NRT1.1 influences drought tolerance, we analyzed the performances of Col-0, *chl1-5*, *cpk6* and *cpk6*/*chl1-5* mutant plants (Fig. S3). The plants of Col-0, *chl1-5*, *cpk6* and *cpk6*/*chl1-5* grown in HN soil were not watered for 15 days for drought stress. Under normal growth condition, all the tested plants showed no significant difference. When exposed to drought stress, *cpk6*/*chl1-5* double mutant plants exhibited slightly wilted in comparison with Col-0, and recovered the drought-sensitive phenotype of *cpk6* (Fig. 6A). The fresh weight of *cpk6*/*chl1-5* were increased compared to Col-0 under drought stress (Fig. 6B). The MDA level and nitrate content of *cpk6*/*chl1-5* double mutant was much lower than Col-0 and *cpk6* mutant plants (Figure 6C-6D). These results totally showed that *CPK6* mutation in *Arabidopsis* decreases drought tolerance in an NRT1.1-dependent manner.

**Figure 6.**
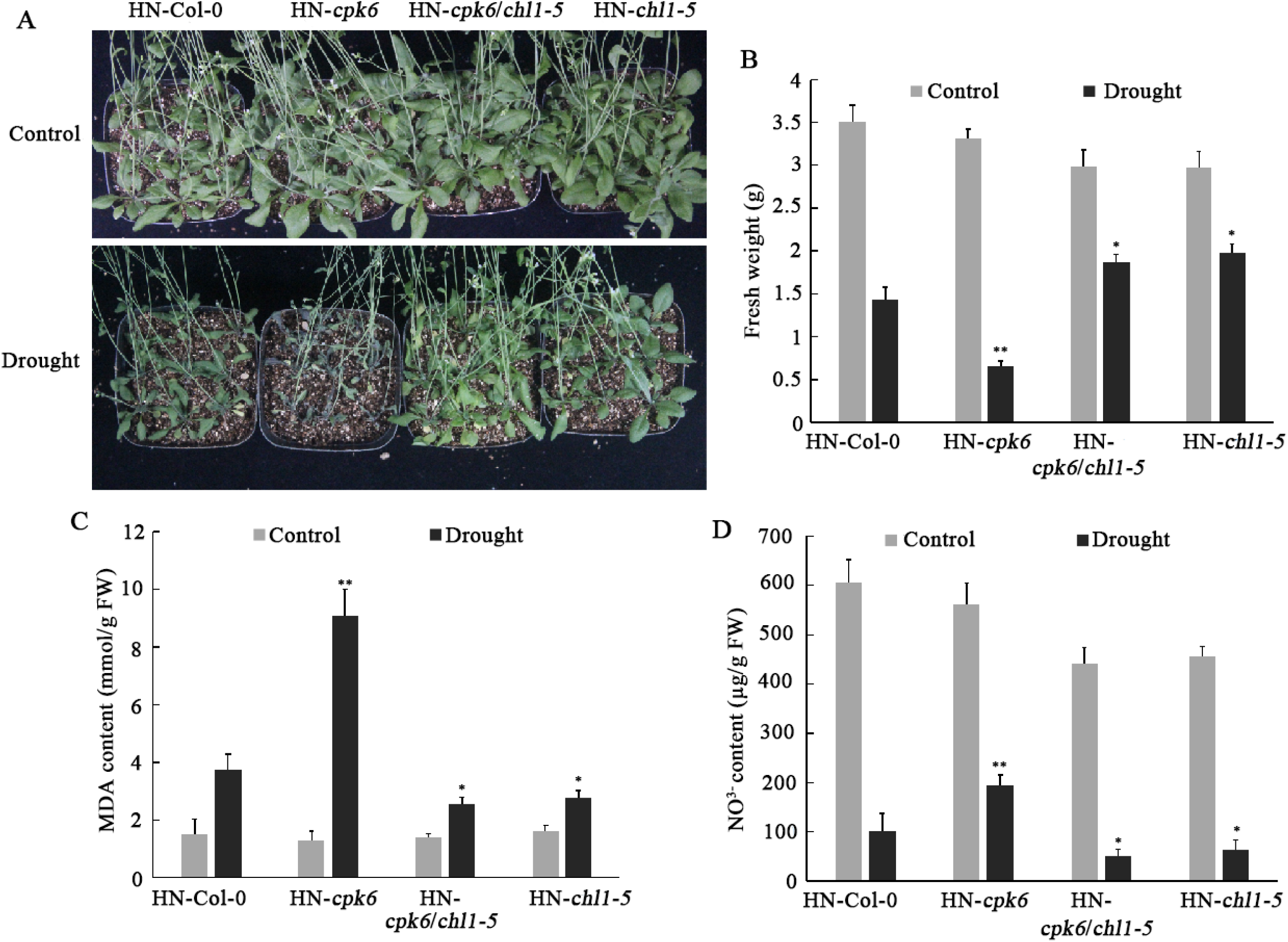
The *Arabidopsis* mutants of *chl1-5* and *cpk6*/*chl1-5* confer drought tolerant under HN condition. **(A)** The drought tolerance phenotypes of the mutant *chl1-5*, *cpk6*, *cpk6/chl1-5* and Col-0. These plants grown in HN soil were treated with drought stress for 15 days and then photographs were taken. **(B)** and **(C)** The fresh weight and MDA contents of the mutant *chl1-5, cpk6, cpk6/chl1-5* and Col-0 plants grown in HN soil were stressed with drought for 15 days. Each column represents the mean±SD. of three independent replicates. **(D)** NO_3_^−^ contents of the mutant *chl1-5, cpk6, cpk6/chl1-5* and Col-0 plants grown in HN soil were stressed with drought for 15 days. All the experiments were performed for three times.

## Discussion

### CPK6 decreases nitrogen absorption of *Arabidopsis* under drought stress by phosphorylating NRT1.1

Drought is an adverse environmental stress resulting from physiological water deficiency as well as the difficulties in nitrogen absorption and utilization in plants. Nitrate metabolism (absorption and utilization) and water deficiency are correlated physiological processes in plants. Upon drought stress challenge, plants need to adjust their physiological states to reduce water transpiration and regulate nutrient absorptions such as nitrogen uptake. CPK6 was characterized in regulating stomatal aperture through phosphorylating and activating the S-type anion channel SLOW ANION CHANNEL-ASSOCIATED 1 (SLAC1) (Mori *et al.*, 2006; Brandt *et al.*, 2012; Scherzer *et al.*, 2012). Considering that SLAC1 and SLAH2 and SLAH3 belong to the same clade of NO_3_^−^/Cl^−^ selective channel families, we thus asked the questions: how does CPK6 link the signals of drought and N metabolism?

To address this question, we firstly compared the phenotypes of Col-0 and the *cpk6* mutant grown in the soil of LN and HN respectively. As our results indicated, the loss function of *cpk6* mutant conferred sensitive to drought stress only under HN condition (Fig. 1). Combining with the results of increased total N content in *cpk6* plants (Fig. 2), we deduce that CPK6 negatively regulates nitrate transporting, and then affects drought tolerance in *Arabidopsis* under HN condition. Protein interaction analysis showed that CPK6 interacts with and phosphorylates the nitrogen transporter NRT1.1 (Fig. 4). Compared with wild type, the phosphorylation level of NRT1.1 in *cpk6* mutant was decreased. Moreover, the phosphorylation level of NRT1.1 was significantly less when the *cpk6* mutant plants grown in the HN soil were treated with drought stress (Fig. 5). These results totally indicate that CPK6 limits N transporting of *Arabidopsis* under HN through upregulating the phosphorylation level of NRT1.1.

### The T^517^ phosphorylation of NRT1.1 influences the drought tolerance of *Arabidopsis*

Phosphorylation analysis showed that CPK6 phosphorylates the residue T^517^ of NRT1.1 (Fig. 5). Knocked out *CPK6* expression resulted from the non-phosphorylated NRT1.1 in T^517^ site, and the lower phosphorylation level of NRT1.1 in the *cpk6* mutant under drought stress (Fig. 5). Given that the higher content of NO_3_^−^ in *cpk6* plants in comparison with Col-0, we deduce that the T^517^ phosphorylation negatively regulates the transporting activity of NRT1.1 in *Arabidopsis* under HN and drought stress.

The T^517^ residue is a novel phosphorylation site of NRT1.1. The mutation T^517^ led to a completely loss of phosphorylation for NRT1.1 protein (Fig. 5D), indicating it is a single and specific phosphorylation site in NRT1.1 in *Arabidopsis* under HN and drought stress. The T^101^ in NRT1.1 is another key site phosphorylated by CIPK23 under LN determining the N-high/low affinity transition (Ho *et al.*, 2009). The *cipk23* mutant was identified to be drought tolerant in comparison with Col-0 (Cheong *et al.*, 2007; Sadhukhan *et al.*, 2019), showing the opposite phenotype to the *cpk6* mutant (Fig.6) (Zhang *et al.*, 2022). These results together suggest that CPK6 and CIPK23 don’t phosphorylate NRT1.1 simultaneously under HN and drought stress. NRT1.1 phosphorylations are respectively triggered by drought stress and environmental N changes, suggesting the universality that multiple-sites of the same protein can be phosphorylated by more than one kinases in response to different stresses (Wang *et al.*, 2020).

Under HN (5mM NO_3_^−^) condition, NRT1.1 can sense N signalling and activates a phospholipase C activity for increasing cytosolic Ca^2+^ levels; The increase of cytosolic Ca^2+^ levels is abolished in *nrt1.1* mutant (*chl1-9*) (Riveras *et al.*, 2015). Consistent with NRT1.1’s role in Ca^2+^ signalling, *chl1-5* and *cpk6/chl1-5* are resilient to drought (Fig. 6). It is also noteworthy that CPK6, serving as an important calcium sensors in plants, mediates ABA signaling and drought tolerance by activating ABF3 and ABI5 transcriptions (Zhang *et al.*, 2020). These results further indicated that the interaction module of CPK6 and NRT1.1 in *Arabidopsis* under drought stress regulates downstream gene expressions probably through the second messenger Ca^2+^ (Zhao *et al.*, 2018; Xiao *et al.*, 2022). Hence, the future works are to characterize the downstream genes and to analysis how they regulate their expressions under HN and drought stress.

## Acknowledgments

We thank many colleagues who have contributed to the researches of promoter analysis and synthetic promoter designing in plants and apologize to those whose works were not cited owing to space limitations. This work was funded by Key Research and Development Projects (No.2018YFA0901000, 2018YFA0901003), China Postdoctoral Science Foundation (2021M692103) and the BIO-Agri. project of SJTU.

## Author contributions

Zuo K.J. designed and conducted the experiments. Ma Q.J. Zhao C.Y. and Hu S. performed the experiments. Zuo K.J. and Ma Q.J. wrote and revised the paper.

